# Characterization of methylation status of the nuclear hormone receptor *DAX-1* in human cancer

**DOI:** 10.1101/2023.11.21.568180

**Authors:** Caroline P. Riedstra, Michael B. Heskett, Christina Tzagarakis-Foster

## Abstract

The orphan receptor DAX-1 plays an essential role in human development, steroid hormone synthesis and the maintenance of embryonic stem cell pluripotency. Recent studies have demonstrated DAX-1 is involved in cancer development, and, depending on the specific cancer type, has a negative or positive effect on cancer growth. In order to better understand the mechanism of *DAX-1* gene regulation in various cancer cell lines, the epigenetic regulation of *DAX-1* was investigated. Following confirming levels of DAX-1 expression at both the mRNA and protein levels, the overall methylation status of the *DAX-1* gene was probed using methylation-sensitive restriction enzyme analysis. To determine the molecular mechanism of DNA methylation of the *DAX-1* gene, chromatin immunoprecipitation assays identified key methylating proteins that localize to specific CpG islands in the *DAX-1* promoter. In conclusion, this study demonstrates that methylation of key cytosine residues in CpG islands within the *DAX-1* promoter play a central role in regulating *DAX-1* expression and varying degrees of methylation result in differences in *DAX-1* expression in human cancer cell lines.

## Introduction

Dosage sensitive adrenal hypoplasia congenita on the X chromosome gene 1 (DAX-1, NR0B1) is an orphan nuclear hormone receptor (NHR) implicated in development and disease. Specifically identified in dosage-sensitive sex-reversal (DSS) and adrenal hypoplasia congenita (AHC), alterations in the levels of DAX-1 expression or mutations in the *DAX-1* gene can lead to critical errors during development (1–4). Knockdown of Dax-1 in mouse embryonic stem (ES) cells results in differentiation, demonstrating the contribution of Dax-1 in maintaining pluripotency in mouse ES cells (5). Although the importance of DAX-1 in mediating AHC and DSS as well as regulating the pluripotent state in mouse ES cells is fairly well studied, the regulation of DAX-1 expression and its mechanism of action still remains elusive largely due to its orphan status. As an orphan, DAX-1 does not bind any known ligand, a key difference compared to other NHRs such as estrogen receptors and androgen receptor (2,6). Additionally, DAX-1 is unique in its ability to repress gene expression by binding to other NHRs. This is due to its uncommon structure at the amino terminus that contains a lysine-rich DNA binding domain (Figure 1a), allowing strong protein-protein interactions (1,6). Within the context of disease, regulation and repression of NHRs can either help suppress cancer, or promote cancer cell growth. Misregulation of gene expression, either upregulation or repression, is common in cancer cells, and epigenetic control plays a key role in tumorigenesis. Transcriptional repression is frequently correlated to increased DNA methylation, thus, tracing methylation status of candidate genes in cancer has proved to be a powerful avenue in cancer research (7). For example, methylation of the Rb1 gene has been investigated as a potential biomarker in bladder cancer progression and development (7,8). Additionally, the epigenetic regulation via hypermethylation of the secreted frizzled receptor proteins (SFRP) family has been implicated in hepatomas and ovarian, colorectal, gastric cancer (9–13). The specific methylation of cytosine residues in CpG islands and the activity of nuclear hormone receptors is a current area of investigation for cancer screening and therapeutic targets (7,8,13–15). Therefore, understanding the epigenetic regulation of NHRs that are clinically relevant, such as DAX-1, could elucidate disease progression and misregulation during growth and development.

**Figure 1.**
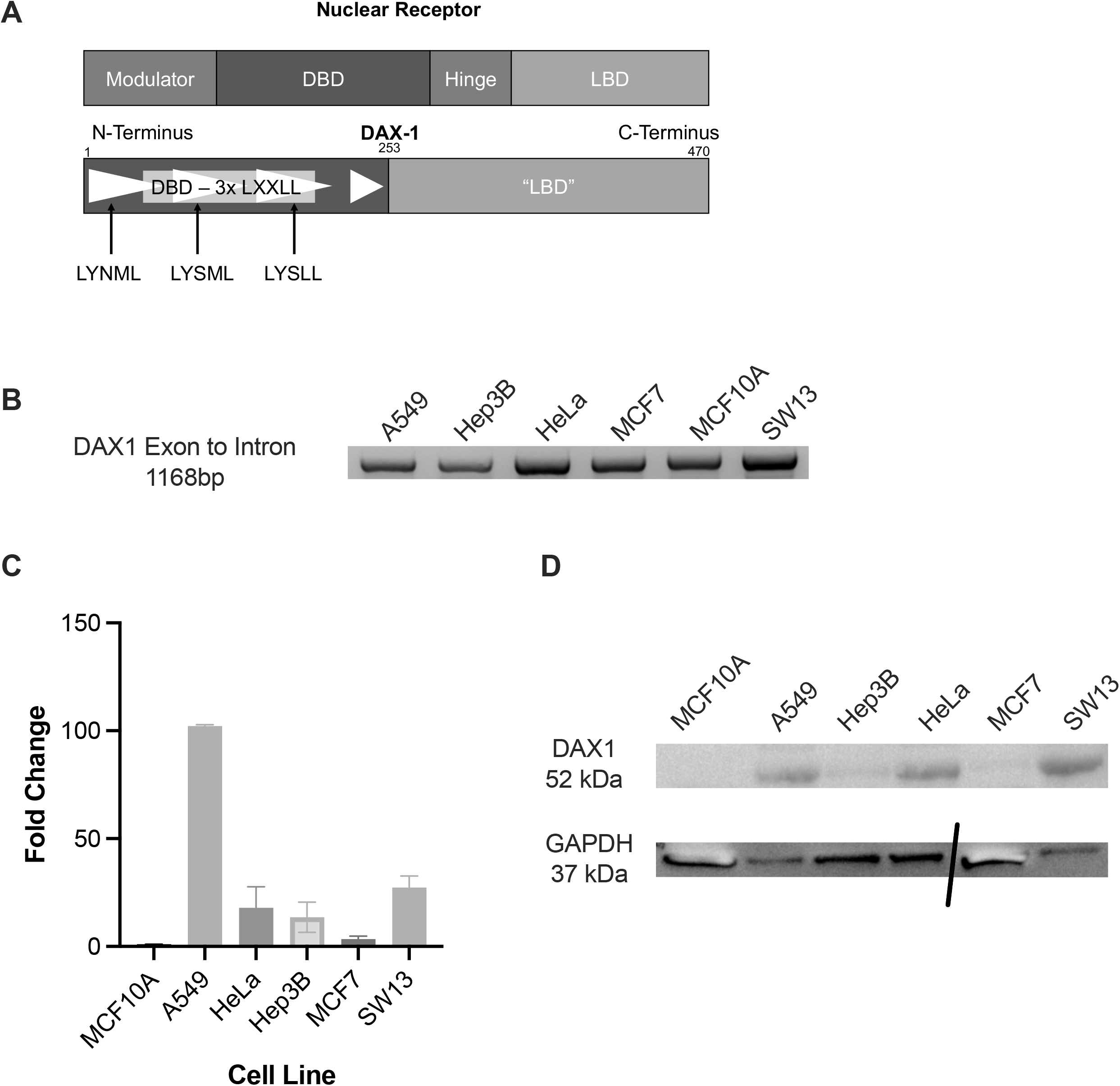
Orphan Nuclear Hormone Receptor DAX-1 in Human Cancer Cells. Unlike other nuclear receptors, DAX-1 does not have a modulator or hinge region and the DNA binding domain (DBD) contains three leucine rich LXXLL repeats that have been shown to mediate cofactor binding in other nuclear hormone receptors (A). MCF10A non-cancerous immortalized breast cells, A549 lung carcinoma, HeLa cervical cancer, Hep-3B liver carcinoma, MCF7 breast cancer, and SW13 adrenal cancer all have DAX-1 detected at the genomic level. (B) Representative qPCR showing that there are varying levels of RNA expression of DAX-1, A549 cells and SW13 cells have the high level of expression when normalized against GAPDH across three replicates (C). Representative western blot to assess protein level expression of DAX-1, break in the gel to show that a lane was skipped while loading (D).

In binding to promoter regions and initiating transcription based on extracellular signaling, NHRs play a critical role in gene expression that is manipulated in cancer, such as the silencing of tumor suppressors and aberrant proliferation (16–18). Previous research identified thyroid hormone nuclear receptors (TR) as regulators of cancer cell growth and correlated diminished expression of TR with the incidence of colon cancer (19). Other NHRs, such as orphans in the subfamily NR4A1/Nur77/NGFIB and the NR4A2 family, have been implicated in the progression of cervical cancer and could therefore be used as therapeutic targets (18). In contrast, repression of the NR4A family in Acute Myeloid Leukemia (AML) was shown to promote tumor suppression (16). Thus, differential expression of NHR families varies between cancer types and plays diverse roles in cancer progression and repression. The differential expression is, in part, due to epigenetic alterations such as methylation. One significant example is the hypermethylation of NR0B2 or small heterodimer partner (SHP). SHP is a tumor suppressor, however, in hepatocellular carcinoma hypermethylation in the promoter region of the SHP gene leads to epigenetic silencing (17). Evidence of epigenetic regulation of other nuclear hormone receptors playing key roles in cancer provides a strong basis for the investigation of oncogenic correlations of differential expression DAX-1 in cancerous versus healthy tissue.

DNA methylation occurs at the C5 position of cytosine residues and can be carried out by a variety of different methylation modifying proteins, ultimately leading to the repression of gene expression (20–28). Different methylating proteins include the diverse family of DNA methyltransferases (DNMTs), methyl-CpG binding domain proteins, and Kaiso and Kaiso-like proteins. Each of these families consist of numerous proteins that can methylate different cytosines under various conditions, making methylation a complex system of repression in different locations and to diverse extents. Critically, DNA methylation is an epigenetic mark that has been studied in human disease, X-chromosome inactivation, and genomic imprinting disorders (7,24,29–34). With a specific focus on cancer, methylation status has been used as a biomarker for cancer screening as well as an indicator of prognosis. For example, methylation of the Rb1 gene in bladder cancer and hypermethylation of DNA mismatch repair genes hMLH1 and hMSH2 in ovarian cancer have been identified as potential prognostic markers (13,15). Additionally, hypermethylation and the resulting repression of genes in the SFRP family have been implicated in hepatomas, gastric cancers, and colorectal cancers (9–12). Derived from such studies, the importance of methylation in cancer identification and progression emphasizes the value of continuing such research.

Studies investigating the methylation status on a large scale have identified key patterns in the degree of methylation (hyper, hypo, and standard) and location of impactful methylation. This initiative, titled the Human Epigenome (24,35), is an expansive effort as hundreds of alterations in methylation status can occur simultaneously across a multitude of tissues. Thus, narrowing in on the diseased state provides clarity to specific methylation occurrences. Such screens helped identify twenty-two methylation markers in eight genes where hypermethylation is significant in patient-derived lung carcinoma. Notably, these statistically significant sites of differential methylation in tumor versus healthy tissue could be categorized into functional categories of proteins: transcription factors, growth regulation and signaling, metabolic function and nonspecific. Regardless of the role methylation status plays in tumor progression, a consistent status of hypermethylation in cancerous tissues can be used as a biological marker not specific to cancer stage (7,36). Furthermore, comparison of the frequency of hypermethylation sites in colorectal cancer cells versus healthy tissue, highlights CpG islands as common targets (14).

Cumulatively, the data supporting the investigation of the differential expression of the orphan NHR DAX-1 across various cancer tissues and the epigenetic regulation of its expression is important to further the understanding of both the role of DAX-1 in disease as well as in normal physiological function (17,37–40). The results discussed below examine *DAX-1* expression in mammary gland tissue, metastatic breast cancer, lung carcinoma, cervical carcinoma, hepatocellular carcinoma, liver carcinoma, and adrenal carcinoma. Particular emphasis was placed on deciphering methylation patterns of CpG islands in the promoter region. By honing in on this region and identifying trends in the epigenetic regulation of DAX-1 across cancer types could lead to methylation status as a biomarker for cancer progression.

## Results

### Characterization of DAX-1 in human cell lines

Each of the human cell lines examined have detectable DAX-1 at the genomic level (Figure 1b). Initial examination of mRNA isolated from each cell line indicated that *DAX-1* is differentially expressed across the various cancer tissue samples (Figure 1c). The control non-cancerous mammary gland cells (MCF10A) were found to be notably low expressors of *DAX-1*, which was expected. Mammary gland cells, unlike adrenal cells that require DAX-1 to regulate steroid hormone production, can vary a great deal in their production of nuclear hormone receptors and generally do not express significant levels of DAX-1 (41). When normalized against the housekeeping gene, GAPDH, the cell lines examined fall into two groups: those with relatively high expression of *DAX-1* and those with negligible amounts (Figure 1c). Those that have higher levels of *DAX-1* expression were found to be the lung carcinoma (A549) and adrenocortical carcinoma (SW-13) cell lines. In comparison, breast cancer of the mammary glands (MCF7), cervical carcinoma (HeLa), and hepatocellular carcinoma (Hep-3B) were found to be relatively low expressors. To further investigate the variable levels of DAX-1 expression, protein expression was analyzed across the different cell lines (Figure 1d). Consistent levels of expression between RNA and protein was seen in the two groups of high expressors (HeLa, SW-13, and A549) and low expressors of DAX-1 (MCF7 and Hep-3b). However, a slight difference in the DAX-1 expression between the RNA and protein was detected in the MCF10A. This could be attributed to translational regulation, specifically when observing a change such as the shift from average RNA expression of DAX-1 in the MCF-10A cell to an undetectable amount at the protein. These initial experiments highlight the potential for the regulation of *DAX-1* at the transcription level that varies between cancer types.

### Identification of targeted regions of methylation

DNA methylation is known to be a common mechanism to switch genes into an “off” position and studies have shown that methylation near gene promoters correlates with low or no expression (26). The 5’ region of DAX-1 upstream of the transcriptional start site and into the first exon were found to contain a CG rich region containing many CpG sites, with total CG content between ∼60-80% (Figure 2a). Downstream of the first 1500 base pairs of the DAX-1 gene, the CG content drops substantially. To investigate the methylation status of the promoter region of the *DAX-1* gene, a three-step approach was taken examining specific cancer types of interest. Previous research in the nuclear hormone receptor field served as a guide for selecting the cell types to investigate. The MCF7 epithelial cancer cell line, derived from a breast adenocarcinoma, and the MCF10A breast epithelial cell line were examined in order to compare cancer versus normal immortalized cells. Additionally, the cervical (HeLa) and liver (Hep3B) carcinoma cells lines have previously been shown to express *DAX-1* (40,42). Also included were lung (A549) and adrenal (SW13) carcinoma cell lines that were anticipated to have varying levels of *DAX-1* expression based on their roles, or lack of, in hormone regulation. CpG islands, particularly those of 5’ – CCGG – 3’ sequence, are frequent targets of cytosine methylation and subsequent repression of gene expression. Thus, eight CpG islands in the DAX-1 promoter region and first exon, identified from the MethPrimer analysis, were crudely examined for methylation status using methylation specific restriction enzyme analysis.

**Figure 2.**
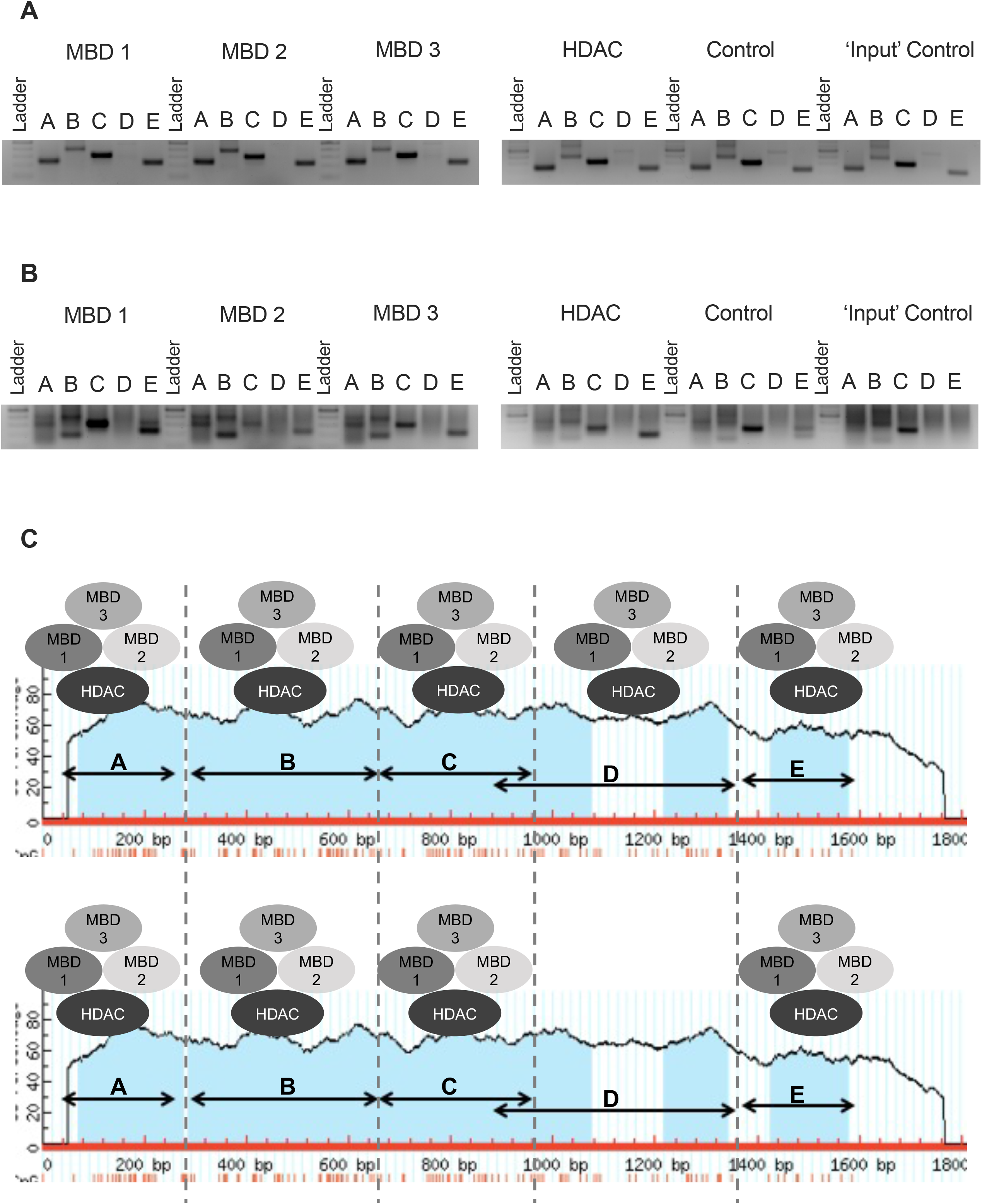
Methylation status of DAX-1 regions following methylation specific restriction enzymes Hpa-II and MSP-1. From the Li Lab MethPrimer (43) software showing CpG rich regions in the DAX-1 promoter and the regions isolated using spcific primers (A). Intensity results quantified using densitometry on representative DNA gels following digests (B-F).

The *DAX-1* promoter was segmented into five regions to increase the specificity of the analysis, thereby narrowing each amplified region into sequences containing three or fewer unique CpG islands. Additionally, by using two isoschizomer enzymes in tandem, the results of this survey could provide more specific information in regards to the location of the methyl group(s) and degree of methylation (Figures 2b-f). From this assay, three cell types were selected for further investigation, as well as the identification of a single CpG island of interest based on its location and the variability of methylation across all cell lines. Continuing with the noncancerous control (MCF-10A), which showed average levels of DAX-1 mRNA expression and low protein expression, this cell line had the highest variability in methylation status across the CpG islands, with most indicating a hemi methylated or methylated state. The lung carcinoma (A549), a strong expressor of DAX-1 at both the RNA and protein levels, was unmethylated at all eight CpG islands based on the MSRE analysis. Also selected for continued analysis was the breast cancer (MCF7) cell line because of the variability in methylation across all MSRE targeted CpG islands and low expression of DAX-1 at both RNA and protein levels (Figures 1b,c). Since the highest variability of methylation status between the cell lines was observed in the amplified region furthest upstream preceding the transcriptional start site and TATA, a more detailed analysis was taken at this single CpG island in the three cell lines previously identified

To gain a more robust understanding of methylation status between the breast cancer and control mammary gland cells, it was necessary to decipher which methyl binding proteins are present at the CpG islands identified in the DAX-1 promoter. Chromatin immunoprecipitation of three methyl binding domain family proteins (MBD1, 2a, and 3) at the five CpG-rich regions in the DAX-1 promoter highlighted a key difference between the cancerous and control tissues (Figure 3a,b). Primarily, the region directly following the transcription start site that contains three CpG islands is occupied by all three MBDs in the breast cancer tissue, while left entirely unoccupied in the control sample. Critically, methylation in this region could indicate the difference in RNA expression between these two cell lines previously observed (Figure 1c). Since the remaining regions of the DAX-1 promoter are occupied by MBDs in both cell lines, it is likely that at least one other CpG island is partially repressing gene expression as well. Further exploration of alternative CpG islands will be the focus of future studies in our lab (Figure 3c).

**Figure 3.**
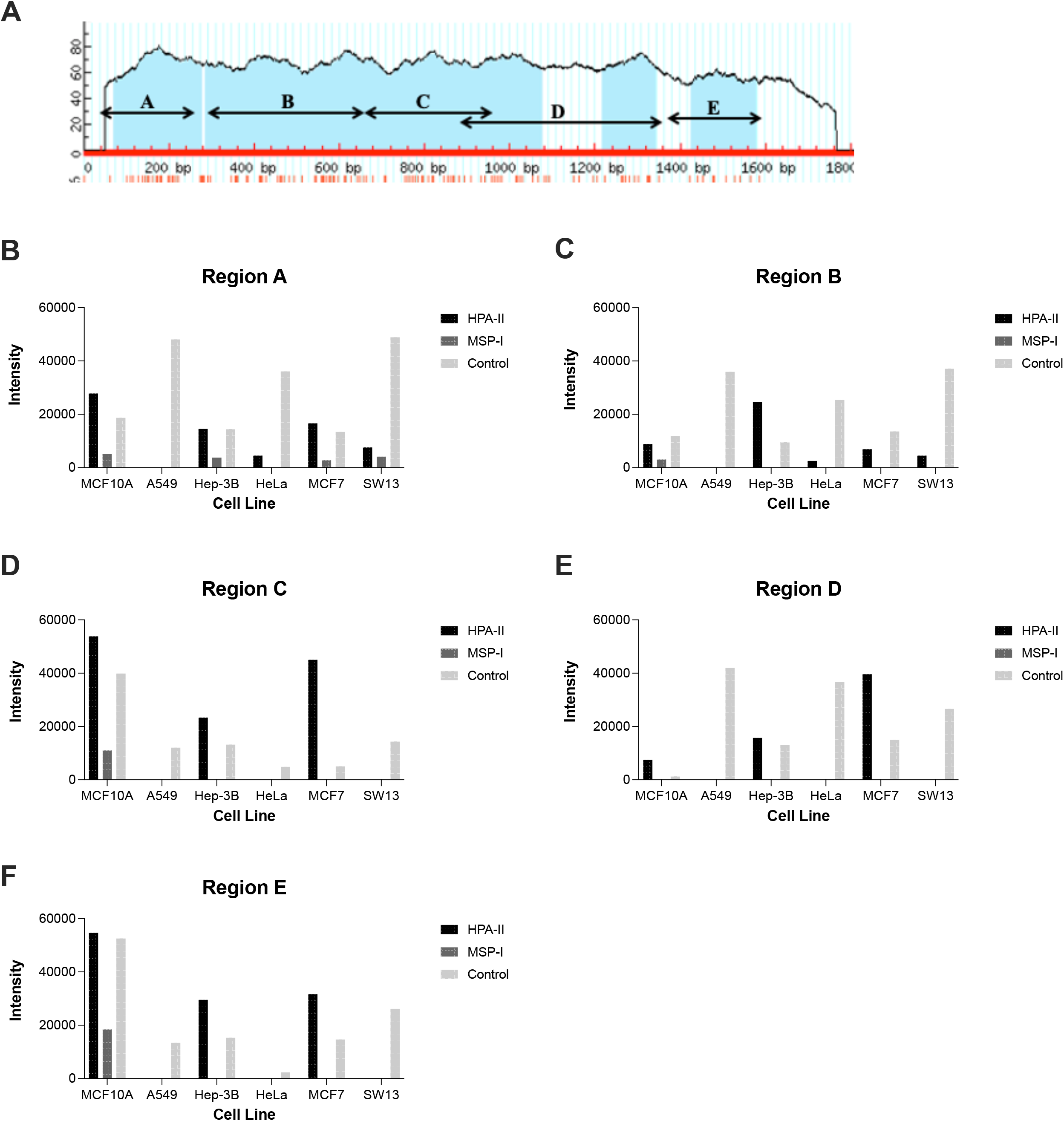
Chromatin immunoprecipitation to identify regions of methyl binding proteins. DNA gels of CHIP results, the presence of a band signifies occupancy of a methyl binding domain protein in the promoter region of DAX-1 isolated from control MCF10A cells (A) and MCF7 cells (B). Summary schematic of CHIP results (C).

## Discussion

The orphan hormone nuclear receptor DAX-1 has been shown to play a critical role in sex determination during developing as well as a suppressor of tumor progression in cancer metastasis. In a survey of different types of metastatic cancer, the experiments in this paper sought to identify trends in *DAX-1* gene expression, as well as means of epigenetic repression. Gene and protein level expression, though qPCR and western blotting, confirmed differential expression of DAX-1 across the cancer types and directed focus towards transcriptional regulation over translational. Through a refined focus on the CpG islands in the promoter region, methylation specific restriction enzyme analysis identified differential methylation between cancer types and within the nine CpG islands in the promoter. To further refine the precise location of the methylated CpG islands, bisulfite sequencing can be carried out on the CpG island nearest the *DAX-1* transcriptional start site in order to confirm that cell lines expressing more DAX-1 are less methylated at this region. In an analysis of methyl binding protein occupancy of the DAX-1 promoter, ChIP experiments highlighted a second region of interest slightly downstream of the transcriptional start site. Ultimately, these results indicate that the methylation status of CpG islands close to the transcriptional start site of the DAX-1 promoter may account for differential expression of DAX-1 in metastatic cancer. Further research in this area could elucidate the degree to which methylation and epigenetic regulation is playing a role in cancer progression. Continuing this line of research into clinical samples of hormone regulated cancers, particularly in the breast, may lead to a biomarker of cancer progression.

## Materials and Methods

### Cell Culture

Human cell lines were obtained from American Type Culture Collection (Manassas, VA). MCF-10A human mammary gland cells were used as the non-tumorigenic control and were cultured in MEBM basal medium supplemented with components included in the MEGM^TM^ Mammary Epithelial Cell Growth Medium SingleQuots^TM^ Kit (Lonza, Hayward, CA). A549 human lung carcinoma cells were cultured in Dulbecco’s Modified Eagles Medium (DMEM) (Invitrogen, Carlsbad, CA) supplemented with 2mM L-Glutamine (Gibco, Waltham, MA), 10% heat inactivated and charcoal stripped Fetal Bovine Serum (FBS) (Cytiva Life Sciences, Marlborough, MA) and antibiotics (Gibco). HeLa human cervical adenocarcinoma cells, Hep-3B human hepatocellular carcinoma cells and MCF7 human mammary carcinoma cells were cultured in Eagle’s Minimum Essential Medium (EMEM) (Gibco) supplemented with 10% heat-inactivated and charcoal stripped fetal calf serum and antibiotics. SW13 human adrenocortical carcinoma cells were cultured in high glucose DMEM (Invitrogen) supplemented with 10% fetal calf serum and antibiotics. Cell cultures were incubated at 37°C in a humidified 5% CO_2_ incubator.

### Prediction of DAX-1 CpG Island

DAX-1 genomic DNA sequence (ID:190) from *Homo sapiens* was obtained from the National Center for Biotechnology Information’s (NCBI) GenBank database. Calculation and visualization of a CG rich region in the promoter region of the human DAX-1 genomic DNA sequence was accomplished with MethPrimer software, using default prediction parameters. CpG islands were defined as a DNA segment with a minimum length of 100bp, >50% CG content, and an observed-to-expected CpG ratio greater than 60%. Observed-to-expected CpG ratio is calculated where:

observed = (number of CpGs/length of sequence),

expected = (((number of C + number of G) / length of sequence) / 2)2.

### Western Blot

Cells were harvested and whole cell lysates were prepared using the Cell Extraction Buffer kit (Invitrogen) and Halt^Tm^ Protease Inhibitor Single-Use Cocktail (Thermo Scientific). Equal amounts of protein (10µg) were separated on 4-15% Bis-Tris SDS-PAGE (Invitrogen) and transferred to a PVDF membrane. Following transfer, PVDF membranes were washed in 1X TBS-Tween and blocked in 5% BLOTTO (prepared in 1X TBS-Tween) for 60 minutes at room temperature. Membranes were incubated overnight at 4°C with anti-NR0B1 (Abcam Ltd, Cambridge UK) or anti-GAPDH (GeneTex) antibodies. After washing in 1X TBS-Tween, membranes were incubated with the appropriate HRP-labeled secondary antibody (BD Pharmingen) diluted 1:2000 in 5% BLOTTO in 1X TBS-Tween for 60 minutes at room temperature. Immunoreactive bands were visualized using the chemiluminescent SuperSignal^TM^ West Femto Maximum Sensitivity Substrate (Thermo Scientific) according to the manufacture’s protocol. Blots were imaged and analyzed on the BioRad ChemiDoc XRS system (BioRad, Hercules, CA).

### RNA, cDNA and qPCR analysis

To determine the level of DAX-1 expression in the different cell lines, RNA was extracted using the Monarch® Total RNA Miniprep Kit (New England Bioloabs, Ipswich, MA) with DNase I treatment to remove any genomic DNA contaminants. Reverse transcription was performed using the Maxima H Minus First Strand cDNA Synthesis Kit (Thermo Fisher) according to the manufacturer’s directions. Quantitative real-time PCR (qPCR) was performed using PerfeCT(T) SYBR® Green Fast Mix (QuantaBio) and qPCR amplification and analysis was performed using the BioRad CFX-96 system. All of the primers used are listed in Table 1.

**Table 1.**
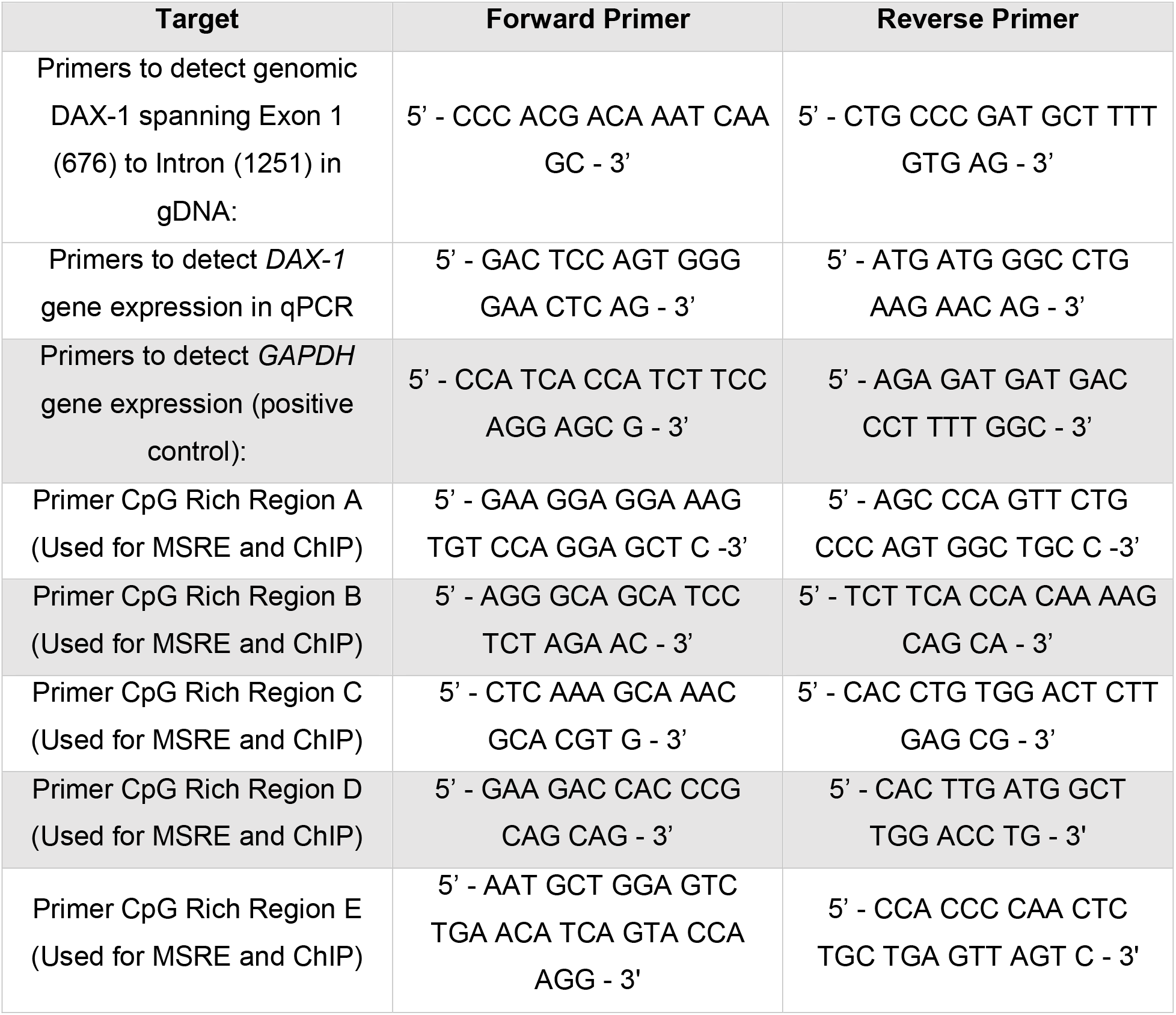
Summary table of the primers used in PCR and qPCR.

The relative expression levels of DAX-1 was determined by the 2^−△△Ct^ method after normalization to the endogenous gene GAPDH. Experiments were performed in triplicate and four independent experiments are summarized.

### MSRE

Regions of interest were determined based on the frequency of CpG (5’ – CG – 3’ and 5’ – CCGG – 3’) islands in the DAX-1 sequence using the Li Lab MethPrimer program (43). Corresponding primers were designed with Primer3 software (44–46). Methylation Specific Restriction Enzyme (MSRE) analysis was carried out using genomic DNA (gDNA) isolated from cell pellets of confluent T75 flasks using the PureLink® Genomic DNA Mini Kit (Invitrogen). Equivalent amounts of gDNA from the different cells lines was digested in a 37°C water bath for four hours with *Hpa*II and *Msp*I isochizomer enzymes (NEB, Inc.) Following digestion, products were purified with the Monarch PCR DNA cleanup kit (NEB, Inc.) and analyzed by endpoint PCR. Quantitative analysis of PCR products was done by densitometry and carried out using ImageLab software (Image Lab Software, Life Science Research, Bio-Rad).

### Chromatin Immunoprecipitation

Cells for chromatin immunoprecipitation (ChIP) were grown to confluence on two 10 cm plates. Cells were collected with trypsin and the ab500-ChIP kit (Abcam) protocol was followed based on cell number. Cross-linked and control non-crosslinked chromatin was fragmented via sonication with a Misonix S-4000 sonicator for 7 minutes at 30 second intervals while on ice. In addition to the non-crosslinked DNA control, a second control using the anti-HDAC3 antibody (rabbit polyclonal, abCAM) was used. For the chromatin immunoprecipitations, 3µg of the following: anti-MBD1 antibody (rabbit, polyclonal, abCam catalog# ab2846-100), anti-MBD2a antibody (rabbit polyclonal, abCam catalog# ab3754-100), and anti-MBD3 antibody (rabbit polyclonal, abCam catalog# ab3755-100). Sheared chromatin that did not undergo any antibody binding was included as an input control. Reactions of sheared chromatin and respective antibodies were incubated with rotation overnight at 4°C and purified the next day with a 50% slurry of protein A agarose beads and manufacture recommended wash steps. Isolated products were electrophoresed through at 2% agarose gel stained with ethidium bromide and imaged on the GelDoc imager and camera system (BioRad, Hercules, CA).

## Acknowledgments

We would like to acknowledge the contributions of James Sikes, PhD in assistance with software analysis of sequencing results. We would also like to acknowledge Dishaa Ramesh and Katerina Fargas for contribution to cell culture.

## Notes

### Competing Interest Statement

The authors have declared no competing interest.

